# RNAinsecta: A tool for prediction of pre-microRNA in insects using machine learning algorithms

**DOI:** 10.1101/2022.03.31.486617

**Authors:** Adhiraj Nath, Utpal Bora

## Abstract

Pre-MicroRNAs are the hairpin loops which produces microRNAs that negatively regulate gene expression in several organisms. In insects, microRNAs participate in several biological processes including metamorphosis, reproduction, immune response, etc. Numerous tools have been designed in recent years to predict pre-microRNA using binary machine learning classifiers where predictive models are trained with true and pseudo pre-microRNA hairpin loops. Currently however, there are no existing tool that is exclusively designed for insect pre-microRNA detection. In this experiment we trained machine learning classifiers such as Random Forest, Support Vector Machine, Logistic Regression and k-Nearest Neighbours to predict pre-microRNA hairpin loops in insects while using Synthetic Minority Over-sampling Technique and Near-Miss to handle the class imbalance. The trained model on Support Vector Machine achieved accuracy of 92.19% while the Random Forest attained an accuracy of 80.28% on our validation dataset. These models are hosted online as web application called RNAinsecta. Further, searching target for the predicted pre-microRNA in insect model organism *Drosophila melanogaster* has been provided in RNAinsecta using miRanda at the backend where experimentally validated genes regulated by microRNA are collected from miRTarBase as target sites. RNAinsecta is freely available at https://rnainsecta.in

## 1. INTRODUCTION

MicroRNAs (miRNA) are a class of non-coding RNA which regulate gene expression. They are typically ~22 bp long and bind to the 3’ untranslated region (3’ UTR) of target mRNAs to induce mRNA degradation and translational repression (1), however, recent studies have suggested that they also bind to the 5’ UTR, coding region and gene promoters (2). It was first discovered in *Caenorhabditis elegans* by Ambros and Ruvkun groups in 1993 (3, 4). Since then it has been discovered in a large number of species across different kingdoms.

MiRNAs are produced in the nucleus by RNA Pol II and III as long primary miRNA (5, 6), and are capped and tailed like mRNAs (7).They are processed into pre-miRNA by Drosha and the stem loop is further processed by Pasha. The pre-mirna is transported to cytosol by Exportin 5 where it’s cleaved by Dicer and loaded into the AGO protein which binds to the mRNA with the help of RNA-induced silencing complex (RISC). MiRNA binds to the mRNA by forming a Watson-Crick basepair at 3’UTR which is known as “seed region”.

The role of miRNA is crucial in insects as it is reported to participate in a wide range of biological activities (8). Changes in the miRNA profiles have been observed during metamorphosis where miR-100/let-7/miR-125 cluster has been found to participate in wing morphogenesis in both hemimetabolan and holometabolan species(9–11). In reproduction, during ovarian development miR-309 plays a critical role in female *A. aegypti* mosquitoes and during spermatogenesis miR-7911c-5p is upregulated in *B. dorsalis*.(12, 13). Several miRNAs has been found to play important role in the regulation of immune related genes(14, 15) and also during insecticide resistance where the genes responsible are downregulated with the help of miRNAs, miR-2b-3p is found to be involved in regulation of metabolic resistance(16, 17).

In recent years numerous tools have been designed to predict pre-miRNA using machine learning approaches by training data to classify pre-mirna hairpin from pseudo pre-miRNA. Tools for species specific pre-miRNA detection like TarMirPred (18) and phylum specific such as ViralMir(19) have also been developed. Most of the tools use the characteristics of the hairpin loop as features for the classification (20–26). Most tools consider 8,494 non-redundant human pseudo hairpins as the negative dataset (20, 21, 27–30), however selection of negative dataset still remains a challenge and careful consideration is required to make efficient binary supervised classification models (31, 32).

Genomic hairpin sequences which are not pre-miRNA viz., mRNA, tRNA and rRNA are also used as negative set (33). However, inclusion of such collection of pseudo-hairpins give rise to the class-imbalance problem. This issue is addressed in tools like HuntMi, where thresholding classifier score function is combined with receiver operating characteristics (ROC)(34), microPred where the concept of undersampling majority class and oversamp ling minority class was used (35) and DeepSOM addresses this issue by creating self-organizing maps (36).

Tools have also been developed to search for potential miRNA target sites in a genomic sequence such as miRanda, Pictar, mirmap (37–39) etc. These tools search for potential target sites for a given sequence in a gene by calculating likelihood, allowing wobble basepairing and reward and complementarity at 5’ end.

As miRNA plays a major role in insects and yet a tool which is exclusively dedicated for its detection is not available, we have designed an ML based pre-miRNA prediction tool while handling the class imbalance problem using SMOTE. The pre-miRNA predicted as positive can also be used to search for probable targets in genes of *Drosophila melanogaster* chromosomes that have been reported to be regulated by microRNAs. The tool is available at https://www.rnainsecta.in with a user-friendly interface and the source code is available at GitHub.

## 2. Methods

### 2.1. Data Preprocessing

#### Dataset

We downloaded all the available insect pre-miRNA sequences from miRBase (40). A total of 3855 sequences were collected and labelled as positive set for the ML classification. For negative dataset we used the 8494 pseudo-hairpin sequence along with genomic sequences of different insects containing hairpins of length below 250 bp. We downloaded 20,654 PCGs from the GenBank and after filtering 14,802 sequences were added to the negative dataset which made it a total of 23,269 sequences.

#### Handling Class Imbalance

As there is a huge difference in ratio of positive to negative classes, state of the art techniques were implemented to address class imbalance. Two strategies namely Synthetic Minority Over-sampling Technique (SMOTE) (44) and Near-Miss (NM) (45, 46) were used to balance the dataset. Packages in python are available for implementation of both techniques (47).

### 2.2. Features for the classification

We calculated different measurable properties of both the classes based on which the ML models were trained. A total of 93 features were calculated as described below:

#### Triplet Element scores

We calculated the minimum free energy (MFE) and secondary structure was using RNAfold of ViennaRNA package (41). We used the standard notation for RNA secondary structure containing dot and bracket where each nucleotide was marked either a dot ‘.’ or bracket ‘(‘ & ‘)’ corresponding to its unpaired and paired state respectively. Left bracket ‘(‘ indicates paired nucleotide at 5’ end and has the complementary base ‘)’ located at 3’ end of the given sequence (42). We used TripletSVM’s method for calculating the triplet element scores where, given any three adjacent nucleotides, there are eight (2^3^) possible structure compositions: ‘(((’, ‘((.’, ‘(..’, ‘…’, ‘.((’, ‘..(’, ‘.(.’ and ‘(.(’, taking ‘(’ for both instances of paired nucleotide. Considering the middle nucleotide, there are 32 (4 × 8) possible structure-sequence combinations, which are denoted as ‘C(((’, ‘G((.’, etc. (20).

#### Base Composition

The percentage of each nucleotide, i.e. %A, %C, %U and %G, dinucleotide counts and their percentage, eg. %AA, %AG, %CG, etc, base pair composition i.e. %C+G and %A+U using inhouse python script.

#### Structural and thermodynamic features

Number of stems, loops, loop length and number of basepairs were calculated from the secondary structure using regular expression and were used as features. A motif containing more than three contiguous base pairs in the secondary structure is termed as stem. The features dG, dP, dD, dQ, normalized Shannon entropy, MFE1 and MFE2 were adapted from miPred (27). dG is calculated by taking the ratio of MFE to the Length. dP is the Base Propensity which is the ratio between total number of basepairs and length. MFE1 is the ratio between dG and %(C+G) and MFE2 is the ratio between dG and number of stems.

MFE3 and MFE4 features were implemented from microPred (35). MFE3 is the ratio between dG and number of loops while MFE4 is the ratio between dG and the total number of bases. dD is the adjusted basepair distance (43) and zD is normalized dD.

### 2.3. Classification and performance evaluation

There are numerous machine learning algorithms which are used to classify any given dataset based on the features provided that are mapped to a given label. This approach is known as supervised learning and in our experiment positive and negative are the provided labels. Let *m* be training samples as n-dimensional vectors X_i_=[X_i1_,…,X_in_]^T^ such that **L**={(x_i_,y_i_)}; i=1,…,*m*, where x_i_∈R^n^ and y_i_∈{−1,+1} are the response variables, i.e. the labels. Given a dataset for training where each vector belongs to one of the two possible classes, a supervised algorithm constructs a model that can predict whether any new data vector whose class is previously unknown belongs to one class or the other.

#### Classification Algorithms

The training was performed on different ML algorithms viz., Support Vector Machine (SVM), Random Forest (RF), Logistic Regression (LR) and k-Nearest Neighbours (KNN) to classify the positive and negative labelled miRNA from the calculated features. The dataset was divided into a training set (X_train) which consisted of 75% of the data and a testing set which consisted of 25% (X_test) of the data (48).

#### Hyperparameter Tuning

Different parameters of the ML algorithms were applied to the SMOTE and NM balanced datasets to classify the positive and negative microRNA. For SVM, linear and Radial Basis Function (RBF) kernels were used along with different values of the Cost function (C_SVM_ value) and gamma. In the case of RF, different values for the number of trees, learning rate, maximum depth, minimum number of sample split and sample leaf were used. For KNN, number of neighbours and different distance matrices were used and in case of LR, regularization strength (C_LR_ value) along with different solvers were used.

We used python’s scikit-learn package to choose the hyperparameters for training each algorithm (49). Initially, a wide range of hyperparameters were chosen and the classification was done by using a model selection package called RandomizedsearchCV which randomly chooses different parameters to train the ML algorithm with 10-fold cross validation. The parameters were then fine-tuned using GridsearchCV, where the training was performed using each of the possible combinations of the provided parameters along with 10-fold cross validation (CV).

The classifiers with different hyperparameters were chosen based on their 10-fold crossvalidation score and were then modelled with the best performing parameters. X_test was used then used to evaluate their performance.

#### Performance Evaluation

The best performing models for each classifier were selected for both SMOTE and Near-Miss balanced dataset after adjusting the hyperparameters. The Cross-Validation score (CV Score), which is the mean accuracy of the 10 folds, was considered for performance and the parameters yielding the highest mean accuracy were selected for each classifier. The best parameters for each classification algorithm were chosen and the models were evaluated on X_test dataset to check for overfitting during the training. Their performance was calculated based on the following classical classification measures: sensitivity (*SN*), specificity (*SP*), Accuracy (*Acc*) precision (*p*), harmonic mean of sensitivity and precision (*F*_1_) and Matthew’s correlation coefficient (*MCC*):

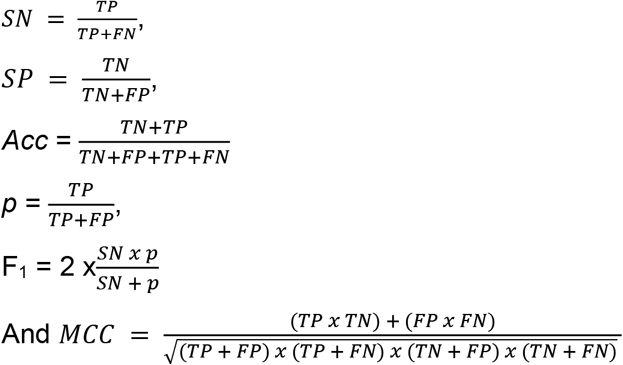

where *TP, TN, FP* and *FN* are the number of true-positive, true-negative, false-positive and false-negative classifications, respectively.

Finally to generalize the model, an independent test dataset was created from the sequences of *Spodoptera frugiperda* (50) and *Tribolium castaneum* (51, 52) which were not used in the modelling and remained unseen to the models were considered as positive dataset and insect CDS of 250bp length fetched from GenBank were considered as negative dataset. A total of 999 sequences were considered as the validation dataset (V_test) of which 464 were positive and 535 were negative.

### 2.4. Web application and Target Searching

#### Web server implementation

The selected trained models were implemented in a backend server using python’s Flask API on a cloud platform along with NGINX as reverse proxy as given in Figure 1. Input from the user is received by NGINX as an HTTP request which it sends to the backend Flask server using reverse proxy. The request from NGINX is interpreted by the Flask API using Gunicorn which is a python WSGI (Web Server Gateway Interface) HTTP server. A port was assigned to the Flask process by Gunicorn with which NGINX communicates. Gunicorn is run in background using Supervisor which also monitors the process, keeps track of errors and restarts the app in case it stops. In the Flask app the sequence features are calculated and the selected ML model predicts if the given sequences are pre-microRNAs or not. The results are sent to the browser from the Flask app as Jinja2 templates. HTML, CSS, Javascript and JQuery were used along with Jinja2 template for the frontend design.

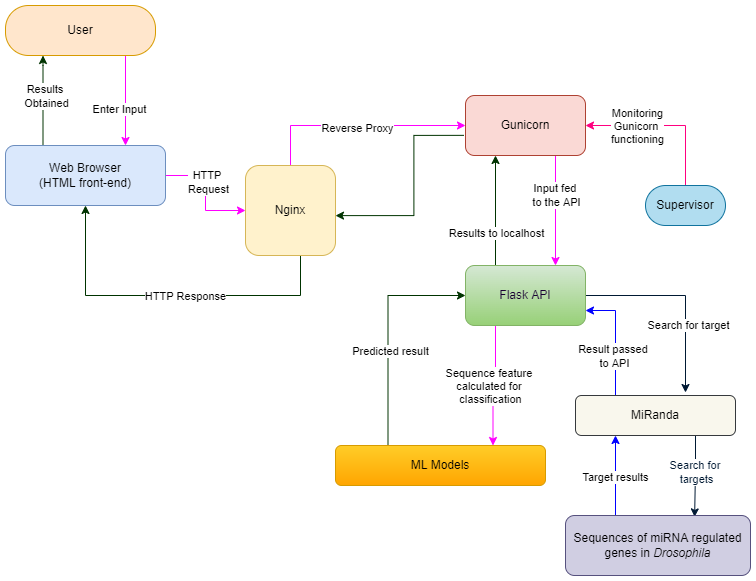

#### Searching for miRNA targets in *Drosophila melanogaster*

For the predicted pre-miRNA users can also search for miRNA targets in Drosophila miRNA regulated genes. Experimentally verified miRNA and their target gene IDs of *Drosophila melanogaster* were fetched from MirTarBase (53). Parent IDs were retrieved from the target gene ID list using eutilities (54), with which the CDS of the genes were downloaded from Flybase (55). MiRanda, which is a popular miRNA target searching tool, was implemented to enable users to search for potential miRNA targets for their pre-miRNA in the CDS of reported genes regulated by miRNA. Users need to choose the directionality of the sequence they want to consider as miRNA from the pre-miRNA after cleavage by Drosha. In case of multiple hairpins users will need to specify which hairpin they want to select as the cleavage site. MiRanda runs in default parameters on the server, for more accurate results users are advised to run it with custom parameters in its standalone version.

## 3. Results

### 3.1. Data Preprocessing

#### Datasets

The number of true microRNA sequences obtained for each species from miRBase is given in Table 1.

**Table 1:** The number of microRNA sequences collected from each species for building the machine learning model.

| Organism | No. of precursors |
| --- | --- |
| *Ixodes scapularis* | 49 |
| *Parasteatoda tepidariorum* | 148 |
| *Rhipicephalus microplus* | 24 |
| *Tetranychus urticae* | 52 |
| *Daphnia pulex* | 44 |
| *Marsupenaeus japonicus* | 5 |
| Triops cancriformis | 148 |
| Aedes aegypti | 155 |
| Anopheles gambiae | 130 |
| Apis mellifera | 254 |
| Acyrtosiphon pisum | 123 |
| Bactrocera dorsalis | 80 |
| Biston betularia | 2 |
| Bombyx mori | 487 |
| Culex quinquefasciatus | 74 |
| Drosophila ananassae | 76 |
| Drosophila erecta | 101 |
| Drosophila grimshawi | 82 |
| Drosophila melanogaster | 258 |
| Drosophila mojavensis | 71 |
| Drosophila persimilis | 75 |
| Drosophila pseudoobscura | 210 |

#### Handling Class Imbalance

The SMOTE dataset consisted of 23,269 instances for both the classes making it a total of 46,538 instances. In case of NM both the classes had 3855 instances making it a total of 7,710 instances.

### 3.2. Feature Selection

The f score and *p*-value of all 91 features are given in Supplementary Material 1. *p*-value of >0.05 was kept as cut-off for selection after which 89 features were finally considered for the training.

### 3.3. Classification and performance evaluation

#### Selection of parameters

The initial parameters were selected based on the best performing models for both SMOTE and NM datasets. Results of the CV score for various classifiers with different parameters are given in Supplementary material 2 for both SMOTE and NM datasets. The overall CV score of NM was found to be lower than the SMOTE dataset. The best parameters for all the classifiers are given in Table 2 which was used in final model preparation and evaluation.

**Table 2:** Sets of best performing parameters obtained after gridsearching through different values for each classification algorithm. Selection was based on the highest 10 fold cross-validation score.

| Classifier | SMOTE Best CV Score | SMOTE Best Parameters | NM Best CV Score | NM Best Parameters |
| --- | --- | --- | --- | --- |
| SVM | 0.99459 | C<sub>SVM</sub> = 10,<br>Kernel =: RBF,<br>Gamma= 0.01 | 0.952789 | C<sub>SVM</sub> = 1,<br>Kernel =: RBF,<br>Gamma= 0.001 |
| Random Forest | 0.98609 | No. estimators= 1600<br>Max depth = 30,<br>Bootstrap = 'False',<br>Min sample leaf = 1,<br>Min sample split = 2, | 0.944617 | No. estimators= 1400<br>Max depth = 30,<br>Bootstrap = 'False',<br>Min sample leaf = 1,<br>Min sample split = 2, |
| Logistic Regression | 0.9627 | C<sub>LR</sub> = 70,<br>Solver = Newton-cg | 0.946952 | C<sub>LR</sub> = 100,<br>Solver = Newton-cg |
| KNN | 0.98199 | Metric Distance: Manhattan,<br>No. of neighbours = 2 | 0.92166 | Metric Distance = Euclidean,<br>No. of neighbours = 7 |

#### Performance Evaluation

The X_test consisted of 988 positive and 5799 negative entries. Each model obtained from SMOTE and NM was tested on the same dataset so that there is no sampling bias in their comparison. The selection of such large datasets for testing gives a better understanding of their performance. Table 3 contains the performance measures for all the selected classifiers.

**Table 3:** Performance measure for each classifier of SMOTE and NM.

| Classifier | Acc | SP | SN | MCC | p | F<sub>1</sub> |
| --- | --- | --- | --- | --- | --- | --- |
| SVM_SMOTE | 0.97451 | 0.975685 | 0.967611 | 0.90525 | 0.871468 | 0.917026 |
| Random Forest_SMOTE | 0.983498 | 0.992757 | 0.92915 | 0.934044 | 0.95625 | 0.942505 |
| Logistic Regression_SMOTE | 0.969501 | 0.971029 | 0.960526 | 0.8882 | 0.849597 | 0.901663 |
| KNN_SMOTE | 0.970679 | 0.983101 | 0.897773 | 0.885446 | 0.900508 | 0.899138 |
| SVM_NM | 0.421836 | 0.331954 | 0.949393 | 0.270949 | 0.194929 | 0.323448 |
| Random Forest_NM | 0.436865 | 0.35075 | 0.942308 | 0.281079 | 0.198254 | 0.327586 |
| Logistic Regression_NM | 0.446589 | 0.361097 | 0.948381 | 0.284791 | 0.201853 | 0.33286 |
| KNN_NM | 0.431266 | 0.349026 | 0.913968 | 0.285773 | 0.193031 | 0.318743 |

#### Comparison with existing tools

The selected SMOTE models were further analysed for their performance in comparison to the already developed tools, viz., miPred, microPred, Triplet-SVM, HuntMi and MiPred on the same validation dataset, V_test, comprising of 464 positive and 536 negative sequences. Table 4 consists of the performance measures for each of the tools.

**Table 4:** Comparative performance analysis between available tools and trained models tested upon independent validation dataset.

| Tool | Acc | SP | SN | MCC | p | F<sub>1</sub> |
| --- | --- | --- | --- | --- | --- | --- |
| Triplet-SVM | 0.754755 | 0.798131 | 0.704741 | 0.625745 | 0.751724 | 0.727475 |
| MiPred | 0.48048 | 0.128972 | 0.885776 | 0.325557 | 0.468643 | 0.612975 |
| miPred | 0.778779 | 0.785047 | 0.771552 | 0.654075 | 0.756871 | 0.764141 |
| microPred | 0.713714 | 0.570093 | 0.87931 | 0.574294 | 0.639498 | 0.740472 |
| HuntMI | 0.618619 | 0.306542 | 0.978448 | 0.414076 | 0.550303 | 0.704422 |
| SVM | 0.921922 | 0.945794 | 0.894397 | 0.854779 | 0.934685 | 0.914097 |
| RF | 0.802803 | 0.695327 | 0.926724 | 0.676982 | 0.725126 | 0.813623 |
| LogR | 0.677678 | 0.450467 | 0.939655 | 0.513201 | 0.59726 | 0.730318 |
| KNN | 0.761762 | 0.583178 | 0.967672 | 0.614122 | 0.668155 | 0.790493 |

#### ROC

The ROC curve (Receiver Operating Characteristic curve) is used to measure the performance of a model at different classification thresholds. It is a plot between True Positive Rate (Sensitivity) and False Positive Rate (1 - Specificity). Higher the AUC (Area Under the Curve) of ROC, the better is the model at classifying, i.e. higher degree of separability. The ROC-AUC of the models is given in Figure 2.

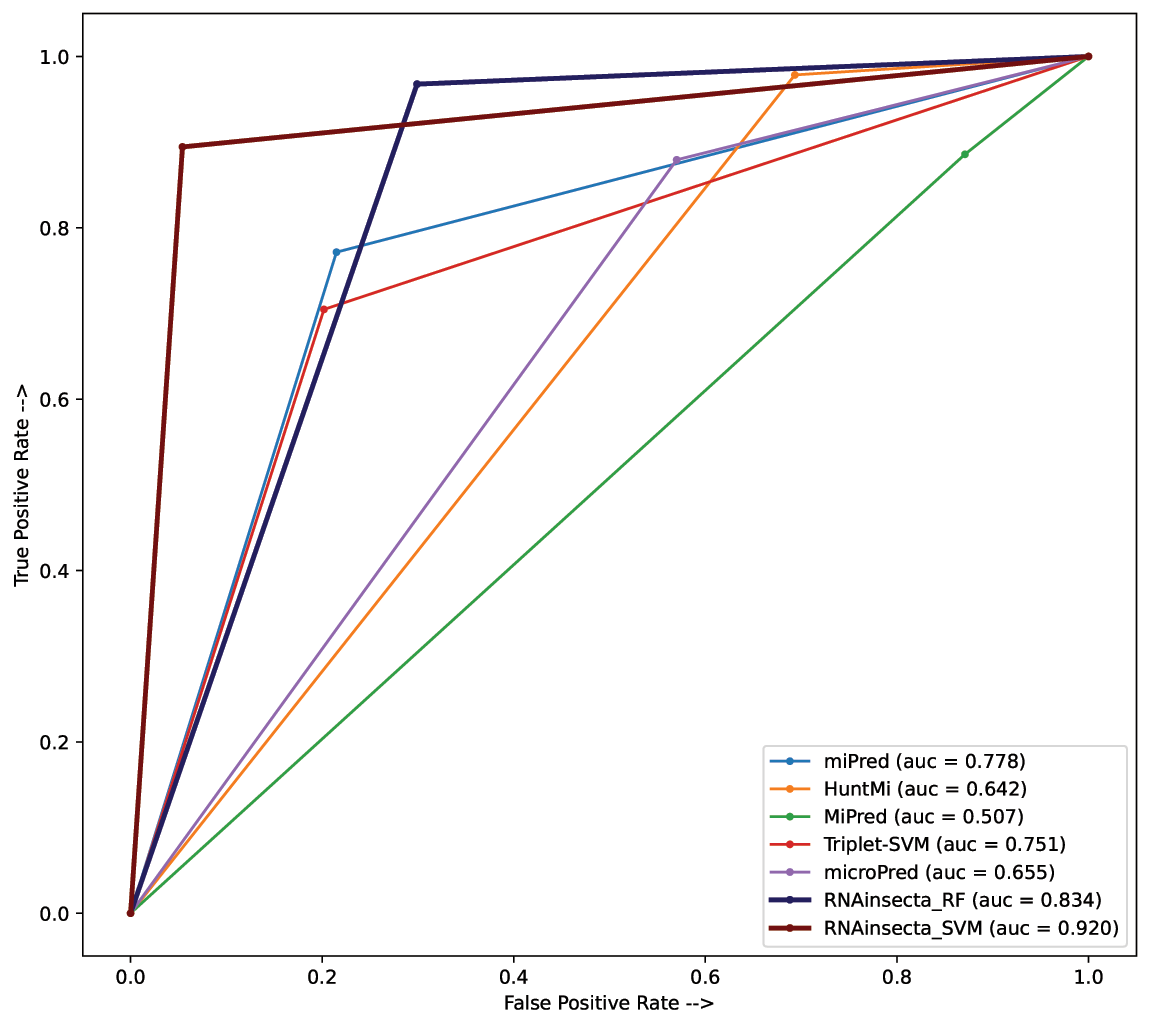

#### Performance on related phyla

The performance of RNAinsecta was measured across species from other phyla and compared with miPred which has so far been better than other tools in our analysis. Their comparative performance is given in Table 5.

**Table 5:**
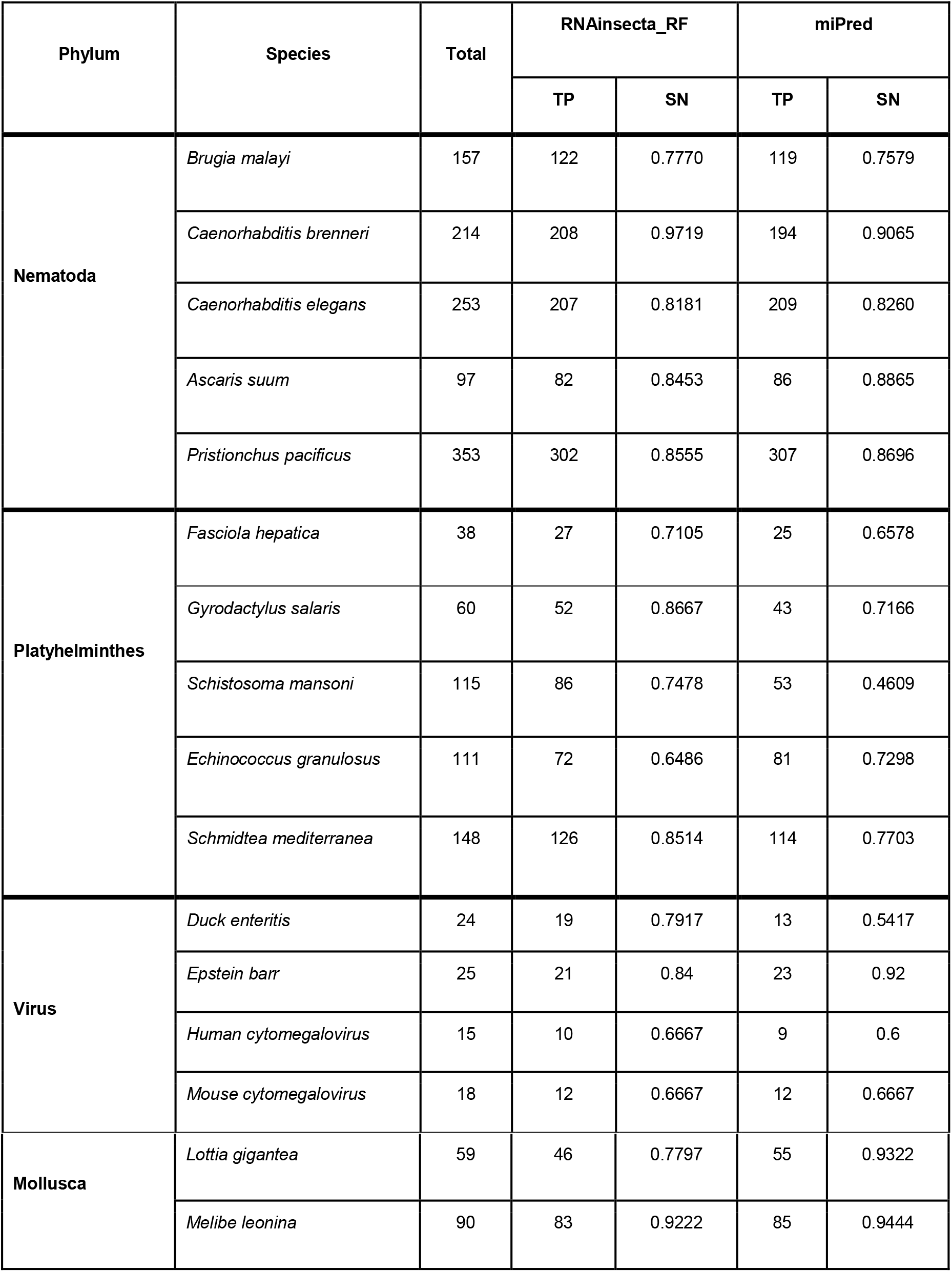
Performance of RNAinsecta in comparison with miPred for prediction of pre-miRNA across related phyla.

### 3.4. Web Application and miRNA Targets

The number of targets for each chromosome of *Drosophila melanogaster* is given Table 6. A total of 174 targets were collected and stored. MiRanda target searching program searches in these targets and displays top the 100 results.

**Table 6:** No. of targets from each chromosome of *Drosophila melanogaster*

| Chromosome | No. of Sequence |
| --- | --- |
| 2L | 34 |
| 2R | 7 |
| 3L | 46 |
| 3R | 55 |
| X | 23 |
| 4 | 9 |

## 4. Discussion

### 4.1. Data Preprocessing

#### Dataset

The longest pre-microRNA reported for insects is 222 bp long produced by *Plutella xylostella* (miRBase ID: MI0027331) for which we kept the length of sequences in negative dataset below 250bp. The consideration of large negative datasets made it a typical imbalanced dataset classification problem since the positive to negative class ratio was approximately 1:6. The problem with such classification is that the majority of data considered for the classification belongs to a single class and hence the results are misleading.

#### Handling Class Imbalance

In SMOTE (synthetic minority oversampling technique), the minority class is oversampled with synthetic data which is extrapolated in such a way that the expectation value of the class remains the same. Near-Miss (NM) approach on the other hand undersamples the majority class by eliminating majority class instances which are close to a minority class instance.

### 4.2. Feature Selection

The features MFE3 and MFE4 had the highest f score value which suggests that the data points for these two features differ the most between positive and negative datasets. The features UU, %AA, G(.( and U.(( had the lowest f score value with p-value above 0.05 suggesting the positive and negative data points do not significantly differ for which they were discarded from the final dataset which brings a total of 89 features for the model training.

### 4.3. Classification and Performance Evaluation

#### Hyperparameter Tuning

Performance of SVM was greatly influenced by the parameter gamma and a value of 0.01 was found to be optimum for this classification. The value of C_SVM_ was assigned to 10 and 1 for SMOTE and NM respectively.

RF classifiers gave best performance with the value of minimum sample leaf and sample split fixed at 1 and 2 respectively while the remaining parameters were further tuned. Maximum depth of 30 was found to be optimal for modelling and hence was assigned in both SMOTE and NM datasets. The number for estimators yielding best performance for SMOTE was 1600 which was slightly more than NM which had 1400 estimators.

Logistic Regression classifiers performed poorly on SAG and SAGA solvers. The performance was slightly better with Liblinear, however the best performance was obtained with Newton-CG solver. CLR value was tuned after fixing the solver and a value of 70 and 100 were found to be optimum for SMOTE and NM respectively.

KNN classifiers used either Manhattan or Euclidean as distance metric. The performance was greatly influenced by the number of neighbours as expected and a number 2 and 7 neighbours were found to be optimum in case of SMOTE and NM respectively.

#### Performance Evaluation

CV score helps in the initial choice of the hyperparameters, however, to regularize the classifiers which performed well, they were tested on X_test that was initially kept separate. In imbalance class testing data, accuracy, sensitivity, specificity, precision and F1 are not the best measures to analyse performance of models as they are not based on the entire confusion matrix. MCC is a better estimator of performance in such cases as it produces a high score only if good results are obtained in all of the four confusion matrix categories (56). The NM models did not perform well on unseen data as was expected from their CV scores. The accuracy of the NM classifiers dropped quite below their CV Scores. As the training data of NM had lesser negative class, the models could not learn to classify non-miRNA which closely resemble true miRNAs and hence suffered from Type I error. The SN of these models were quite high suggesting they learned fairly well to classify positive miRNAs but produced a lot of FP as their SP was low. Hence, these models had poor precision and accuracy for which they were discarded from further analysis.

The SMOTE models performed well on the test data as the models learned to correctly classify non-miRNAs which included insect CDS hairpins that closely resembled true miRNAs. The accuracy of Logistic Regression was the lowest among all the SMOTE models but still had higher MCC and F1 than the KNN model. The MCC of SVM and RF were the highest among all the models.

The performance of SMOTE was better than NM suggesting that with increase in the amount of training data, the performance of these classifiers improves. Hence, for the validation test the SMOTE models were selected for their performance analysis and comparison with already existing tools.

#### Comparison with existing tools

As the negative dataset resembled closely with true microRNA, most of the tools classified them as true microRNA making the Type I error. HuntMi and MiPred had the least Specificity with 0.31 and 0.13 respectively. HuntMI had an F1 score of 70.44% but precision was 55.03%. microPred although had 71.37% accuracy, the specificity was 57%. Triplet-SVM and miPred performed well with MCC of 62.57% and 65.4% respectively, which was the highest among the previously developed tools considered for this experiment. The SMOTE models had fairly good sensitivity but the Logistic Regression and KNN model suffered from the same Type I error. The RF model had accuracy and precision of 80.28% and 81.36% which was higher than all the previous tools. However, the best performance was given by the SVM model with specificity of 94.58% which was the highest among all models used in the experiment indicating it had the least Type I error. The accuracy, precision and F1 score of the SVM model was also highest with 92.19%, 93.47% and 91.41% respectively. However, to achieve such low FP the model was allowed to make few Type II errors for which sensitivity of the model was lower than RF but yet was more than Triplet-SVM and miPred. The MCC score of SVM was 85.48% which was the highest by quite a margin. As RF and SVM models performed better than all the models, both were considered for implementation in a web server called RNAinsecta and the choice of model to select will depend on the users and their experiment.

#### ROC

Triplet-SVM and miPred have lesser FPR than RNAinsecta RF model but more than RNAinsecta SVM. Tools like microPred and HuntMi although have high TPR also have high FPR for which their AUC is less. RNAinsecta SVM and RF had the highest AUC with 0.92 and 0.83 respectively followed by miPred and Triplet-SVM with 0.78 and 0.75 respectively.

#### Performance on other Phyla

RNAinsecta RF performed well on Nematoda with highest prediction specificity. The performance on Platyhelminthes was better as compared to miPred. The performance on Virus was almost same as miPred whereas in case of Mollusc, miPred performed better.

### 4.4. RNAinsecta webserver

The user-interface (UI) has both batch and single sequence searching modes. The batch mode allows maximum 200 FASTA sequences as input. User can choose either SVM or RF model as predicting models. After initial screening, user may check the single sequence search mode for further searching the targets. User can select the orientation of the miRNA to be cleaved from pre-miRNA to be either 3’ or 5’. In case of more than one hairpin, user can choose the cleavage site based on the secondary structure. Finally, the chromosome of *Drosophila* has to be selected where the target has to be searched.

The resulting window will contain the list of possible targets genes and their FlyBase ID along with the miRBase ID of the miRNA that has been reported to regulate that gene.

## 5. Conclusion

In this work we present a new web-based tool for predicting pre-miRNA in insects. We used 87 features from previously reported tools such as Triplet-SVM, miPred, MiPred, etc, relating to the sequence, structural and thermodynamic characteristics of pre-miRNA. These features were trained on various ML algorithms such as SVM, Random Forest, Logistic Regression and KNN for binary classification of true and pseudo pre-miRNA. SMOTE and Near-Miss were used to handle the imbalance in the class, along with 10-fold cross-validation. Two models were selected upon their performance evaluation based on SVM and RF with accuracy of 92.19% and 80.28% respectively, tested on independent validation dataset along with other previous tools.

Further, the target for candidate miRNA produced from the pre-miRNA can be searched for the known miRNA regulated genes in *Drosophila melanogaster*. The target regions are the genes which are known to be regulated by miRNAs and therefore the user can check the details about the gene from the provided hyperlink.

To our knowledge this is the first tool which provides prediction of insect pre-miRNA as well as searching target for the resulting miRNA. The prediction of pre-miRNA in transcriptome from RNA-Seq data can be implemented in the future.

